# Modelling and assessment of combining gilt vaccination, vector control and pig herd management to control Japanese Encephalitis virus transmission in Southeast Asia

**DOI:** 10.1101/430231

**Authors:** Alpha Oumar Diallo, Véronique Chevalier, Julien Cappelle, Raphael Duboz, Fontenille Didier, Durand Benoit

**Author notes:** These authors contributed equally (Alpha Oumar Diallo, Véronique Chevalier).

## Abstract

Despite existence of human vaccines, Japanese Encephalitis (JE) remains a prominent public health problem in Southeast Asia (SEA). JE is caused by a Flavivirus which is transmitted between pigs, the main amplifying hosts, by *Culex* mosquito bites. Therefore, sow vaccination, pig herd management and vector control –or a combination of these three potential control measures, might constitute additional control measures contributing to reduce JE health impact in humans, and economic losses in pig farms. We built a deterministic metapopulation model, combining a pig and a *Culex* mosquito vector population, to represent JE virus (JEV) transmission dynamic within a pig herd. The dynamic of the epidemiological systems resulted from an infectious process, operating in continuous time, combined with the pig breeding process that was modeled based on discrete events occurring instantaneously. We used this model to simulate JEV transmission within a continuum of plausible pig breeding systems encountered in SEA, ranging from backyards to semi-commercial systems. We then analyzed the joint effects of the three tested control measures, namely sow vaccination, pig herd management and vector control, on several indicators characterizing (i) the ability of different pig breeding systems to be simultaneously profitable and allow JEV eradication in the herd, (ii) the impact of JE on pig production and the profitability of gilt vaccination, and (iii) the risk for human beings living in the vicinity of pig herds and/or near pig slaughterhouses. According to our model, herd management has no effect on JEV circulation. Vector control alone is a major control tool but shows paradoxical effects that should be considered in any mosquito based control strategy. Combining sow vaccination and vector control could be an alternative or an additional measure to human vaccination to efficiently reduce both JE incidence in humans and the economic impact of JE infection on pig farms.

**Author summary:** Japanese Encephalitis (JE) still has an important impact on human health in Southeast Asia. Human vaccination is an efficient tool to protect humans but it may not be effective against emerging strains, and poor or remote population may not be able to afford it. Severe outbreaks still occur. JE virus (JEV) is primarily transmitted between pigs and mosquitoes. When infected after sexual maturity, pigs show reproduction disorders leading to economic losses. We propose a modelling approach to investigate the joint effect of three additional control measures, namely sow vaccination, vector control, and pig herd management on JEV transmission dynamic, risk for humans and pigs, and pig breeding sustainability. According to our results, vector control, associated or not with sow vaccination, may be an efficient tool to reduce JE incidence in both human and pigs.

## Introduction

Despite vaccination, Japanese Encephalitis (JE) remains the most important cause of human encephalitis in several Asian countries and the Pacific (1–4). JE is caused by a Flavivirus (Flaviviridae family) (5) which is transmitted between pigs, the main amplifying hosts, by mosquito bites, especially *Culex tritaeniorhynchus* (6, 7). Although JE is known as vector-borne disease, direct transmission may occur between pigs (8, 9). It is admitted that Ardeid birds such as Egrets and Herons are the natural maintenance reservoir (10–12). However other animal species may play a role in the maintenance of the virus (13). In particular, domestic birds may also play a role in transmission processes: an experimental study demonstrated that young ducks and chicken produce a sufficient viremia to infect mosquitoes when biting (14) and preliminary serological results obtained from chickens sampled in Cambodia (P. Dussart, pers. com; S1 Letter) show that they are exposed and produce antibodies against JE. Humans and horses are dead-end hosts (1–4). JE is therefore predominantly a rural disease, even if recent studies showed that peri-urban and urban areas are also affected (15–17).

Each year, JE causes around 68,000 human cases (18). Clinical encephalitis occurs in 1/50 to 1/1000 cases, mostly in children. Then fatality rate can reach 30%, and 30 to 50% of survivors may remain with definitive neurological or psychiatric sequelae. There is no licensed anti-JE drugs available, and the management of patients remains symptomatic (4). In pigs JE infection is symptomatic only when occurring after sexual maturity (after the age of 8 months). In this case the infection can be responsible for reproductive disorder, such as fetal abortion, stillbirth in infected sows and infertility in boars. Infected piglets can also display fatal neurological signs. JE infection in pigs may thus impact breeder livelihood in countries where pig production is the main or one of the main income sources for households (19–21).

To protect humans against JE infection, vaccines have been available since the 1930s and are used internationally (22, 23). Several cost-effectiveness analyses has shown that JE routine or campaign based vaccination reduce JE incidence, especially in children (24–28). An analysis incorporating 14 endemic countries estimated that campaign and routine vaccination would allow to save about USD 19 million in acute hospitalization costs (29). However, human cases still occur.

Recently recommended in temperate or sub-tropical areas to reduce losses due to late infections in sows, pig vaccines, that are commercially available in Asia, may contribute to reduce JE virus (JEV) transmission within pig populations, thus the risk of human infection. These vaccines are widely used in Japan, Taiwan and South Korea (30–32). In Korea, a program of vaccination with a live-attenuated strain [Anyang300] (33) conducted throughout the country for the past 30 years has reduced incidence of disease in swine (34, 35). A modelling survey performed in northern Bangladesh suggested that vaccinating 50% of the total pig population each year would result in an 82% reduction in the annual incidence of JE in pigs (36). A combined use of human and pig vaccination may thus be efficient under given socio-economic environmental and epidemiological conditions to significantly reduce JE impact on public health and pig production (23, 37).

However, both human and pig vaccination have limitations. Human vaccines are expensive, require multiples doses, and remote and/or poor people may not be able to afford it. Some of these human vaccines may not be 100% effective as demonstrated in (38). Additionally, human vaccination alone cannot stop virus circulation, and human cases may re-occur in case of vaccination failure or emergence of a new strain for which current vaccine is not efficient: JE reemerged in South Korea (2010–2015) where vaccination is presumed to have failed to induce lifelong immunity so that older age groups became susceptible again (39). Regarding to pigs, immunity coverage may be hard to sustain due to a rapid turnover of populations and a high unit cost (30). In addition, JEV may still circulate within pig populations, even when vaccinated, as suggested by the results of the vivo challenge performed in Garcia-Nicolas et al (40). However, as a matter of fact, pig vaccination program held in Korea did contribute reducing incidence of disease in pigs but did not prevent outbreaks in the human population in recent years (34). In addition, human and pig vaccines are based on genotype III (GIII) viruses, the dominant circulating genotype in Asia. However there are now several evidence of the replacement of GIII by GI that could negatively modify the effectiveness of current vaccines (41–43). Lastly, the currently available vaccines do not confer full protection against the emerging JEV genotype V strain (44, 45).

Given high and probably under-reported incidence of JE (2), nascent or uncomplete or partially efficient human vaccination programs in some Southeast Asian countries, and poverty that prevent people from being vaccinated or even being informed about JE risk, there is an urgent need to provide recommendations for additional control measures that would significantly reduce JE incidence in Southeast Asia (SEA). As stated in (23), pig vaccination, pig herd management and vector control –or a combination of these three potential control measures, might substantially reduce JEV transmission within pig populations, thus the risk of human disease. In endemic areas, pigs acquire an immunity during the first months of life through colostrum intake and absorption of maternal antibodies (46). They may keep these maternal antibodies up to 3 months (47). Once infected with JEV and seroconverted, pigs remain immune at least three years (48). Since pig’s life expectancy is usually shorter than three years, one can consider pigs as lifelong immunized after JEV infection. Depending on pig herd size and herd management practices, in particular the duration between two successive litters that can be controlled through insemination synchronization, and age at slaughtering, the proportion of immune pigs within a herd may vary and favor or reduce viral circulation between pigs. Vector control can help breaking JE transmission cycle as well, either by targeting adult mosquitoes through insecticide spraying, or juvenile stages with larvacides such as *Bacillus thuringiensis* toxin, extracts of *Piper retrofractum* (Piperaceae) or essential oils as oviposition deterrents (49–52). Use of mosquito nets, either for humans or animals, is also a very effective way to protect them from mosquito bites.

In this work, we analyzed how different combinations of the above-mentioned control measures, i.e. sow vaccination, vector control and pig herd management, could help decrease the risk of JEV transmission to humans and the impact of JE on sows and pig production in rural environments of SEA. To achieve this goal, we built a deterministic metapopulation model that simulated JEV transmission dynamic within a pig herd. We first used this model to compute the yearly limit cycle of JEV transmission dynamics within herds of the two main pig breeding modes encountered in SEA, namely backyards and semi-commercial systems (53). Based on a systematic exploration of the model parameter space, we then simulated the possible combinations of the 3 proposed control measures, to analyze their joint effects on several indicators characterizing: (i) different pig breeding systems that may, at the same time, profitable and to allow JEV eradication in the herd, (ii) the impact of JE on pig production and the profitability of vaccination, and (iii) the risk for human beings living in the vicinity of pig herds and/or near pig slaughterhouses..

## Materials and methods

### General characteristics of pig breeding in Southeast Asia

In most of Southeast Asian countries (Myanmar, Thailand, Laos, Vietnam, Cambodia, Indonesia, Malaysia, Singapore and the Philippines), pig is the most important livestock species and pork is the most preferred meat. Pig rearing is particularly important for smallholders’ livelihood in rural communities, contributing about 20–30% of the rural household income, and even up to 41% in Northern Vietnam (54). Because of industrialization of farming system and transformation of smallholder backyard system to more commercial farming system in response to market demand, pork production in SEA has been growing rapidly over the last decades. However, and except in Thailand where around 80% of pigs produced are from intensive farming systems, smallholders and semi-commercial farms remain predominant, in particular in rural areas. In Vietnam, Laos, The Philippines and Cambodia, about 80% of pigs are raised by smallholders. Most households of rural areas keep less than 5 pigs (55–57), against 5 to 15 in peri-urban areas or in semi-commercial farms (56). In the Philippines and Vietnam, a small farm has less than 20 pigs. In Myanmar, the percentage may go above 90% as commercial pig farming accounts for only a small portion of total pig production. In Cambodia less than 1% of pig producers operate on a commercial level. These farms are well-equipped, and are protected against pathogen introduction (58, 59). Large scale pig farms account for 15–20% of the total SEA pig population. Of these, about 15% belongs to medium scale and 5% belongs to large scale (60).

Regarding backyards and semi-commercial farms, keeping pigs in confinements is common in peri-urban areas, and tends to increase in rural areas. Breeding systems generally comprise three to six pens, which are most likely to be closed by solid walls. Breeders generally own from one to three sows. In “fattener production” one-month-old piglets are purchased from other farms and raised for slaughter. Some households raise only sows that produced piglets that are sold for fattening. Others apply a `farrow-to-finish’ system: piglets born and are raised for slaughter in the same household. Some households applied different combinations of these systems (56). Reproduction is usually not synchronized by farmers in backyard system, where fee based natural breeding is still predominant and payment for boar services is usually in cash or in kind. Oppositely, in semi commercial and commercial farms, synchronization allow breeders to alternate pregnancies and guarantee the simultaneous presence of more than one age group in the farm, and to secure a production flow of piglets throughout the year or at pre-identified periods. Sows usually give birth 2 to 2.5 times a year to piglets which are raised until 6 to 8 months of age and then sent to slaughterhouse.

### Model

The epidemiological system was modeled by a host population (a pig herd) combined with a *Culex* mosquito vector population (mosquitoes). The dynamic of the epidemiological systems resulted from two distinct processes: the infectious process, operating in continuous time was adapted from (61), whereas the pig breeding process was modeled based on discrete events occurring instantaneously. A graphical representation of pig breeding processes and associated infectious dynamics is provided in Fig 1.

**Fig. 1.**
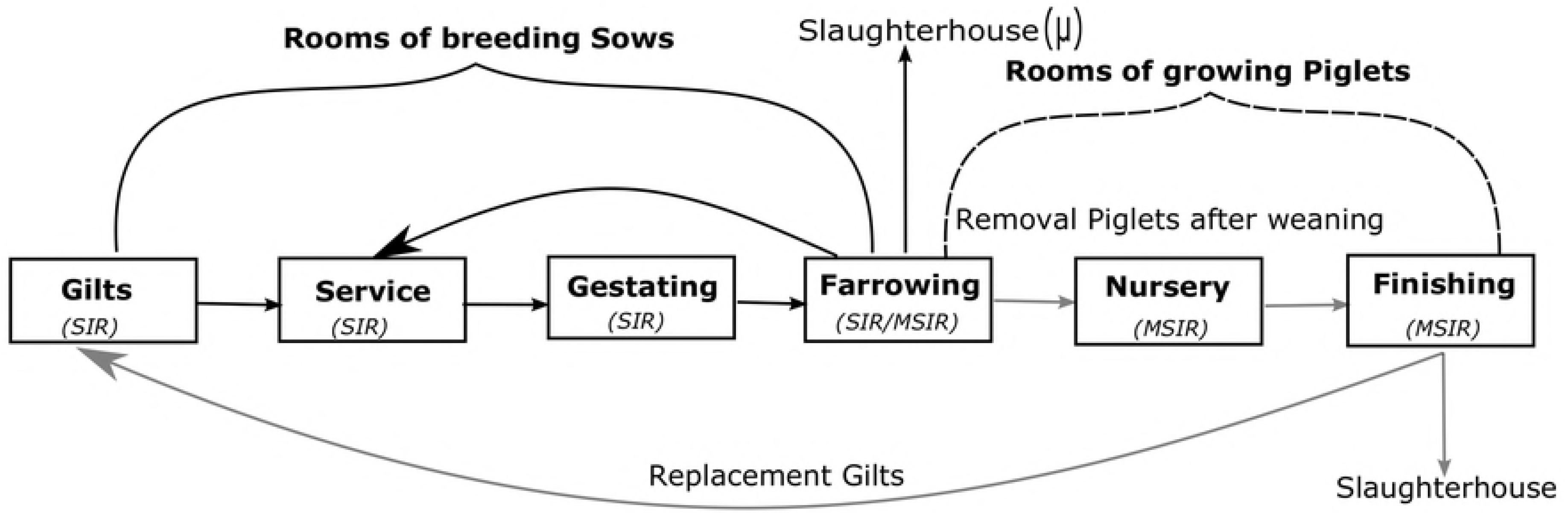
Graphical representation of pig breeding processes and associated infectious dynamics. The white boxes represent the 6 different types of rooms corresponding to each of pig physiological stages. Black and grey solid arrows indicate the breeding sow and growing piglet flows respectively. Black curly brackets indicate rooms where breeding sows are located and black dash curly brackets shows the rooms where piglets are growing. Susceptible-Infectious-Removed (SIR) dynamics are running in Gilt, Service and Gestating rooms, whereas Maternal-Susceptible-Infectious-Removed (MSIR) dynamics are running in Farrowing and Nursery rooms; both SIR and MSIR are running in Finishing room.

#### State variables

The host population represented a pig production unit in which breeding sows give birth to piglets, These piglets grow, are fattened and finally sent to the abattoir. This host population was divided into subgroups of pigs, named hereafter batches, composed of one or several pigs which shared the same production status. Four possible production statuses were distinguished: empty sows, the so-called gilts (*Fe*), gestating sows (*Fg*), aborted sows (*Fa*), and growing pigs (*Pg*), the set of which was denoted *A* = {*Fe,Fg,Fa,Pg*}

Each batch was assumed to be raised in a distinct location which may be a pen or a permanent building (termed below rooms), with possible direct contacts between animals parked in the same room, but not between animals located in distinct rooms. Six types of rooms were distinguished: the gilt room (where nulliparous sows were raised), several service rooms (where sows were placed for insemination), gestating rooms (where pregnant sows were parked), farrowing rooms (where sows gave birth to piglets, and where suckling piglets remain with their mother until weaning), nursery rooms (where piglets were placed after weaning) and finishing rooms (where growing pigs were fattened before being sent to the abattoir). The set of rooms was denoted Z = {z_0_, z_1_,…,z_m_}, where z_0_, was the gilt room.

Five health states were distinguished for hosts *H_h_* = {*M,S,I,R,V*}, with:

– *M*: pigs are protected from JE infection by maternal antibodies,
– *S*: pigs are susceptible to JE infection,
– *I*: pigs are infected and infectious, both for the other pigs located in the same room, and for vectors (*i.e*. hosts are viremic),
– *R*: pigs are immune as a result of natural infection,
– *V*: pigs are immune as a result of vaccination.

The state of the host population was then represented by state variables 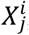, which denoted the number of pigs in the health state *X*(*X ∈ H_h_*), located in the room *i*(*i ∈ Z*), being under production status *j* (*j ∈ A*).

Three health states were distinguished for vectors *H_v_* = {*S,E,I*} with:

– *S*: vectors are susceptible to JE infection and may become infected if they bite a viremic host,
– *E*: vectors are infected but not yet infectious for hosts (extrinsic incubation period),
– *I*: vectors are infectious, and susceptible hosts may become infected if they feed upon them.

The state of the vector population was represented by three state variables *S^v^, E^v^* and *I^v^*, denoting the number of vectors in each health state. Only adult mosquitoes were represented (eggs and larval stages were not taken into account).

#### Infectious process

The infectious process was described by a system of differential equations (Eqs. 1–5 for hosts, and Eqs. 8–10 for vectors), with a daily time step. Piglets protected by maternal antibodies, in *M* state, lose these antibodies at a fixed rate δ. They then become susceptible (*S* state) (Eq. 1). Susceptible animals located in a given room *i* became infected (*I* state) when exposed to infectious bites of vectors (vector-based force of infection (FOI), λ_v_) and/or when exposed to contacts with infectious pigs located in the same room (direct FOI, 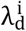) (Eqs. 3a-3c). Infected gestating sows may abort with a probability *α* (Eq. 3c). As in (61), infected pigs were treated as immediately infectious, and we omitted the state “exposed”. Infected (viremic) pigs finally became immune (*R* state) at a fixed rate *γ* (the average duration of viremia was thus1/*γ*). Immunity induced by natural infection (*R* state) of by vaccination (*V* state) was assumed lifelong. 
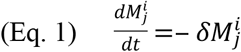

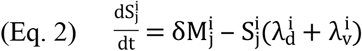

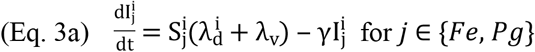

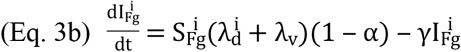

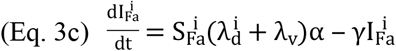

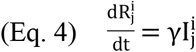

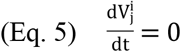

The direct FOI exerted by infectious on susceptible pigs in a given room *i* was assumed frequency-dependent, and depended on a transmission parameter β and on the proportion of infectious pigs (*I* state) among those located in the room *i*: 
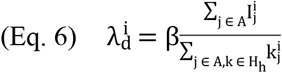

The vector-based FOI exerted by infectious vectors on pigs was identical, whatever the room they were located in. It depended on the proportion of infectious mosquitoes in the vector population, on their biting rate *a*, and on the probability *p* that an infectious mosquito transmits the virus to a susceptible pig when biting. Vector control measures reduced this vector-based FOI according to a parameter *u* representing the efficacy of vector control (0≤*u*≤1) 
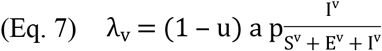

Susceptible vectors (*S* state) became infected and entered the exposed compartment E (infected but not yet infectious) when biting a viremic host (Eq. 8). Infected vectors then became infectious at a fixed rate ε (the average duration of the extrinsic incubation period was thus 1/ε), and remained infectious lifelong (Eqs. 9–10). The renewal of the vector population resulted from a constant mortality rate *μ* (Eqs 8–10), and from the emergence of susceptible mosquitoes (*S* state), as no vertical transmission has been reported in JEV vectors (Eq. 8). The daily number of emergent mosquitoes compensated for the cumulated vector mortality over a yearly cycle, to obtain a stable limit cycle, with an average size of the vector population 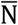 over this cycle. The daily number of emergent mosquitoes could either be constant, or seasonally vary, depending on the time-varying relative emergence level ψ_t_. 
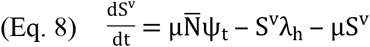

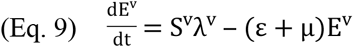

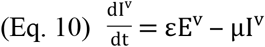

Vectors could fed upon any pig of the herd with the same probability (differences in animal sizes were not taken into account), and the FOI exerted by hosts on vectors (λ^v^) thus depended on the proportion of viremic animals in the herd, on the biting rate *a*, and on the probability *q* for a susceptible vector to become infected when biting a viremic host. Vector control measures reduced this force of infection: 
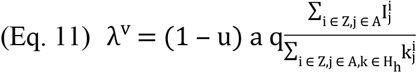

The time-varying relative emergence level ψ_t_ allowed to produce a constant emergence level (and thus a stable vector population size) when η = 0, or, when η = 1 when a yearly sinusoidal cycle peaking at the calendar day ϕ (with 0 ≤ ϕ ≤ 365): 
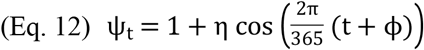

#### Breeding process

The breeding process was described by events occurring instantaneously. Their succession in time represented the reproductive cycle of a sow batch and the growth of their piglets, from birth to slaughter. The reproductive cycle of a sow batch was a (cyclic) sequence of three events: insemination, farrowing and weaning events. The production of finished pigs resulted from the (non-cyclic) succession of four types of events: farrowing, weaning, fattening and slaughter events. The synchronization of batches was controlled by the time interval Δ_Batch_ between successive inseminations of sow batches: 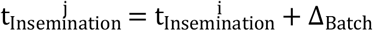 where *i* and *j* were the service rooms where the two successive sow batches were inseminated.

##### Insemination events

Sows located in the service room *i* were inseminated and moved to the first empty gestating room 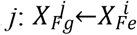 (for each X ∈ H_h_). If 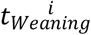 was the time at which this batch of sows had been separated from their piglets and placed into the service room, the insemination event Δ_*Service*_ occurred days later: 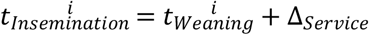

##### Farrowing events

The sows located in the gestating room *i* were moved to the first empty farrowing room *j* (Ev. F1) where they gave birth to their piglet litters of size *L*. Piglets born from immune sows (*R* state) were assumed protected by maternal antibodies (*M* state) whereas other piglets were assumed susceptible (Ev. F2-F3): 
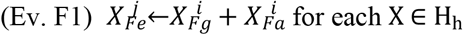

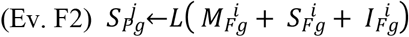

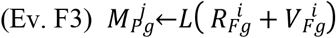

For the batch of sows parked in the gestating room *i* after insemination at time 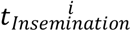 the farrowing event occurred Δ_*Gestation*_ days later: 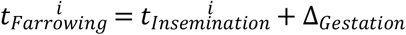

##### Weaning events

At the end of the lactation period, the sows located in the farrowing room *i* were moved to the first empty service room *j* (Ev. W1-W3), whereas their piglets were moved to the first empty nursery room *k* (Ev. W5). Before being moved to the gestating room, some sows were culled with a culling rate *μ*. They were replaced by young females from the gilt room *z_0_*, which could be vaccinated at that time if a vaccination strategy was implemented (Ev. W1-W4). Vaccinating non-susceptible animals was assumed to have no effect on animal health state. 
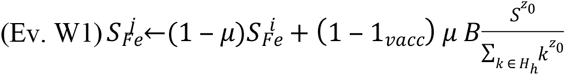

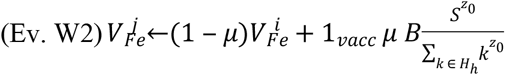
with 1_vacc_ = 1 if vaccination was implemented, and 0 otherwise, and *B* the number of sows in a batch. 
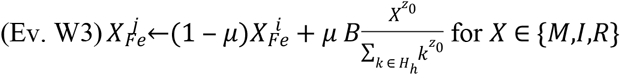

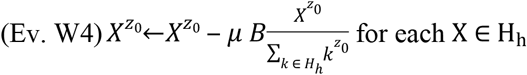

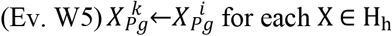

For the batch of sows which had been placed in the farrowing room *i* and had given birth to their piglets at time 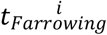, the weaning event occurred Δ_*Laction*_ days later: 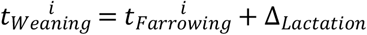

##### Fattening events

The batch of weaned piglets located in the nursery room *i* were moved to the first empty fattening room 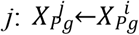 This event occurred Δ_*PostWeaning*_ days after these piglets had been separated from their mother, weaned and placed into the nursery room *i*: 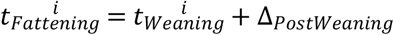

##### Slaughter events

All the finished pigs located in the fattening room *i* were sent to the abattoir, except a fixed number *ρ* of females, kept for sow renewal. These latter were moved to the gilt room *z_0_*: 
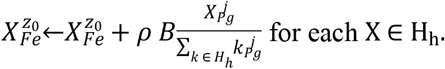
This event occurred Δ_*Finishing*_ days after the pigs had entered the fattening room *i*: 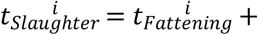 Δ*_Finishing_*

### Initial conditions and parameterization

The average size of the vector population 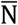 over a yearly cycle was assumed proportional to the total number of pigs in the initial state of the epidemiological system: 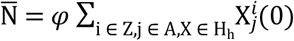, where *φ* is the vector to host ratio, and 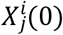 denotes the initial value of the state variable 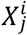 for each X ∈ H_h_. For sow batches (*j* ∈ {*Fe,Fg,Fa*}), 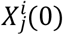, was set to the number *B* of sows in a batch. For growing pig batches (*j = Pg*), it was set to the product of the number of sows per batch by the litter size: 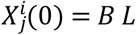. The number of (0) = d batches (sows and growing pigs) as well as, for each batch, the date of the next event of the breeding process, were fixed to satisfy the time interval Δ_Batch_ between two successive inseminations of sow batches, and the above-described event-specific temporal constraints.

A consequence of this parameterization was that, for a given interval Δ_*Batch*_ between sow batches, the total number of hosts (sows and growing pigs) in the initial state of the epidemiological system was proportional to the number of sows per batch *B*. Furthermore, the overall dynamic of the whole epidemiological system was also proportional to the number of sows per batch *B*, as (i) the size of the vector population (averaged over a yearly cycle) was assumed proportional to the initial number of hosts, (ii) the FOI exerted by direct contact between pigs was assumed frequency-dependent, (iii) the FOI exerted by vectors on hosts depended on the proportion of infectious vectors, and (iv) the FOI exerted by hosts on vectors depended on the proportion of viremic hosts. For these reasons, and without loss of generality, the number of sows per batch *B* was set to 1, output variables based on animal counts being computed conditionally to a given size of sow batches, by multiplying the value obtained with *B* = 1 by the specific value of *B*.

The infection was initially seeded assuming that 0.1% of vectors were in *I* state, other mosquitoes being initially in *S* state. All the hosts were initially placed in *S* state.

The parameters of the infectious and breeding processes, as well as the time intervals between breeding events are given in Table 1. When the vector population size showed seasonal variations, the peak of vector abundance was set to the end of July (ϕ = 7 × 30.5).

A time interval of 14 days (or more) with less than one viremic pig (*I* state) in the epidemic dynamic was considered a proxy for epidemic die-out, Indeed, the lifespan of a *Culex* mosquito is between 21 and 30 days (62), and the intrinsic incubation period varies between 7 and 15 days (63, 64). Assuming a newly emerged female would bite a host and get infected the same day, it would become infectious 7 to 15 days later. If it becomes infectious 7 days later, it will remain infectious between 14 days (lifespan of 21 days) and 21 days (lifespan of 30 days). If the newly emerged female becomes infectious 15 days after emergence, it will remain infectious between 6 days (lifespan of 21 days) and 15 days (lifespan of 30 days). The average of the four preceding values (14, 21, 6, 15), 14 days, was considered the average duration of *I* state in vectors. Considering the minor role of direct transmission in disease dynamic (61) a consecutive duration of 14 days with less than 1 viremic pig (*I* state) would result, on average, in the death of all the infectious vectors, without any new infection of susceptible mosquitoes.

**Table 1.** Model parameter: description, values and references.

| Parameters | Description | Value | Values<br>for SA | References |
| --- | --- | --- | --- | --- |
| $\delta$ | Transfer rate from maternally protected to susceptible piglets, per day | 0.03 | NA | (47) |
| $\beta$ | Transmission rate from infectious to susceptible pigs, per day | 0.3 | 0.24;0.3 | (61) |
| $\gamma$ | Pig recovery rate, per day | 0.49 | NA | (3, 65) |
| $\mu_v$ | Mosquito death rate, per day | 0.04 | NA | (5, 66) |
| $a$ | Average daily biting rate per mosquito (average number of bites by mosquito and per pig), per day | 0.25 | 0.2;0.25 | (67) |
| $p$ | Probability that an infectious mosquito transmits JEV to a susceptible pig when biting | 0.11 | 0.09;0.11 | (68) |
| $\epsilon$ | Transfer rate from an exposed mosquito to an infectious mosquito, per day | 0.07 | NA | (3, 63, 64) |
| $q$ | Probability that a susceptible mosquito get infected when biting an infectious pig | 0.37 | 0.3;0.37 | (68) |
| $B$ | Number of sows by batch (sow) | 1 | NA | assumed |
| $L$ | Size of litters (piglet) | 11.5 | 9.2;11.5 | (69, 70) |
| $\mu$ | Sow culling rate | 0.16 | 0.16;0.2 | (69, 70) |
| $\Delta_{\text{Service}}$ | Time spent in service room (day) | 32 | NA | (69, 70) |
| $\Delta_{\text{Gestation}}$ | Time spent in gestation room (day) | 82 | NA | (69, 70) |
| $\Delta_{\text{Lactation}}$ | Time spent in maternity room (day) | 33 | NA | (69, 70) |
| $\Delta_{\text{PostWeaning}}$ | Time spent in nursery room (day) | 49 | NA | (69, 70) |
| $\Delta_{\text{Finishing}}$ | Time spent in finishing room (day) | 105 | NA | (69, 70) |
| $\eta$ | Vector population size | 0;1 | 0;1 | assumed |
| $\Delta_{\text{Batch}}$ | Time intervals between two successive inseminations of sows batches (week) | 3 | NA | assumed |
| **u** | Efficacy of vector control | 0.5 | 0.4;0.5 | assumed |
| **v** | Vaccination | 0;1 | 0;1 | assumed |
| **$\alpha$** | Probability that an infected sow aborts | 0.65 | 0.52;0.65 | (71) |
| **$\phi$** | Ratio vectors per pig | 67 | 67;234 | (61) |

### Model exploitation

#### Transmission dynamics in two Southeast Asian typical breeding systems: village breeding units and semi-industrial farms

The two most common pig breeding systems encountered in Southeast Asia, a village breeding unit and a semi-industrial farm, were modeled based on time interval between Δ_Batch_ successive inseminations of sow batches. A semi-industrial farm represented a pig herd system where sow insemination is synchronized to secure a production flow and sell piglets throughout the year or at pre-identified periods. We used the most common time interval between sow batches, i.e. Δ_*Batch*_ = 21 days (3 weeks) (72).

In a village breeding unit, the first “insemination” of gilts was considered uniformly distributed throughout the year, with Δ_*Batch*_ set to 1 day to mimic real village conditions where pigs are roaming and sows are naturally covered. A village breeding unit represented the set of smallholder backyards of a given village. Each breeder was assumed to own a single sow and raise the piglets and fattening pigs born from this sow. This backyard herd was represented by a batch composed of the unique sow (possibly with its litter), a single batch of weaned piglets and a single batch of finishing pigs, both corresponding to previous litters of the sow. These three batches were assumed to be kept in separate pens (represented by rooms in the model) allowing direct contacts within each pen, but not between pens. A village breeding unit was modeled by a set of such backyard herds, exposed to a unique vector population, without any synchronization of births between sows belonging to different smallholders. We assumed the village large enough to have each day, on average, the birth of one litter of piglets: Δ_*Batch*_ was thus set to 1. Considering the duration of the production cycle of a sow (147 days: see Table 1), this corresponded to a village composed of 147 smallholders.

The limit cycle obtained after stabilization of the epidemiological system was characterized in four situations: a village breeding unit or a semi-industrial farm, in an area where the vector population size was either constant (η = 0) or showed seasonal variations (η = 1).

#### Systematic exploration of parameter space and joint effect of control measures

A systematic exploration of the parameter space was then performed for 3 parameters representing the 3 tested control measures: the vaccination of gilts before their first insemination (1_*vacc*_) with two values: 1 if vaccination was performed, and 0 otherwise; the efficacy of vector control (*u*) with 51 tested values regularly distributed between 0 (no vector control) and 1 (full vector control); and the time interval between insemination of sow batches (Δ_*Batch*_), with 28 values regularly distributed between 1 day and 8 weeks. For each of the 2856 resulting triples, the model was run without (η = 0) or with seasonal variations of vector population size (η = 1). Each model run lasted a simulated period of 12 years, in order to reach the limit cycle, and the last two years of the simulation were used for the computation of five output variables:

– The maximal number of sows compatible with epidemic die-out: this output was chosen as a proxy for epidemic control at the epidemiological unit level (semi-commercial or village). For each possible consecutive period of 14 days, we computed the maximal value of the proportion of viremic hosts. The period of 14 days for which this value was minimal (*q_min_*) was then selected. The maximal number of sows compatible with disease die-out was then 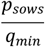 where *p_sows_* was the average proportion of sows among the population of adult pigs during the selected period.
– The abortion rate in gilts: this output was chosen as a proxy for the economic losses induced by JE, and was estimated by the ratio of the total number of abortions to the total count of gilts entering a sow batch for their first insemination. This output variable was only computed when no vaccination was used in gilts (otherwise the number of abortions would by definition be null in the limit cycle)
– The profitability of vaccination: this variable was based on the benefit/cost ratio of vaccinating gilts. The benefit of vaccinating gilts was represented by the product of (i) the total number of avoided abortions *A*, (ii) the litter size *L*, and (iii) the average carcass weight of a finished pig (100 kg), and (iv) the net margin *M* per kg of finished pig carcass. The cost was modeled by the product of the total number of gilts vaccinated *V* by the unitary cost of vaccination *C* (i.e. cost of a vaccine dose and of its administration to an animal). The vaccination was considered profitable if the benefit/cost ratio was greater than one: 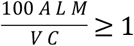, i.e. if 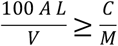. Neither the net margin *M* nor the unitary cost of vaccination *C* could be easily determined and these economic indicators may widely vary according to the country and period. For this reason we analyzed the ratio 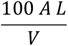, considered as the threshold value of the ratio 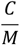 below which vaccinating gilts is profitable: if the number of abortions *A* is very low, the unitary cost of vaccination has to be also very low for the vaccination of gilts to be profitable. Conversely, if the number of abortions *A* is very high, vaccinating sows may become profitable even if the vaccine is expensive.
– The average proportion of infectious mosquitoes: this variable was chosen as a proxy for the exposure to infectious bites of the farmer and his family, as well as the population living around the semi-commercial farm or in the village
– The average proportion of viremic animals among pigs sent to the abattoir: this last output was used as a proxy for the exposure of slaughterhouse staff, and of the population living around the slaughterhouse.

The model was implemented using the R statistical software (R version 3.2.3) (73).

*Sensitivity analysis*

#### Sensitivity Analysis

Sensitivity Analysis (SA) was performed to rank model parameters according to the influence of their value on four of the above output variables: the maximal number of sows compatible with epidemic die-out, the average proportion of abortions among gestating sows, the average proportion of infectious mosquitoes, and the average proportion of viremic animals among pigs sent to the abattoir. SA was separately performed for the two specific epidemiological systems described above: the village breeding unit and the semi-industrial farm. All the model parameters were considered in both SA (except purely biological or zootechnical parameters which were considered fixed) with two values per parameter: the default value and the default value lowered by 20% (Table 1). A two level factorial experimental design was used, resulting in 2^11^ distinct combinations of parameter values. Data were analyzed using a classical analysis of variance (ANOVA) method, taking into account the main effect and first order interactions. Sensitivity indices were then computed, one for each parameter, and one for the interactions between parameter pairs (74). These indices reflected the proportion of variance in the model outputs induced by the variation of the parameter (or parameter pair) values. Parameters were finally ranked from the most influential to the least one. Sensitivity analysis was performed using the R statistical software (R version 3.2.3) with the package “aov” (73).

## Results

### Limit cycle

After stabilization of the epidemiological dynamics, in a village breeding unit located in an area where the vector population size was constant (Fig. 2A), the proportion of viremic host and vectors remained constant, as expected. The proportion of viremic pigs was the lowest in sows (black line) and fattening pigs (purple line): these oldest animals were exposed to infectious bites or to direct transmission for a longer time than piglets. Oppositely, the proportion of viremic animals was the highest after weaning (red line), in young piglets of about one month: most of them had lost their maternal antibodies, as the average duration of the *M* state is approximately one month (Table 1). Due to the protection provided by maternal antibodies, the proportion of weaned piglets (orange line) was slightly lower than that of weaned piglets. The same trends were observed in the semi-industrial farms (Fig. 2B), as well as in both breeding systems with a seasonal vector dynamic (Figs. 2C and 2D). In a semi-industrial farm, the synchronization of sow inseminations induced only small variations of the proportions of infected hosts and vectors (Fig. 2B). Oppositely, the seasonal variations of the vector population size induced marked variations of the proportion of viremic hosts (Figs. 2C and 2D). The proportion of infectious vectors was about 1.5% in a village breeding unit located in an area where the vector population size was constant (Fig. 2A), and varied around this value in a semi-industrial farm (Fig. 2B). For the village breeding unit, and a seasonal vector population dynamics (Fig. 2C), the proportion of viremic sows remains the lowest and almost constant, with a very small peak that corresponds to infection of the yearly renewal sows, the other ones having largely been exposed because of age. There are two peaks of infectious vectors: the first one occurs, by construction at the end of July, calendar day 213. This first peak is concomitant with a global increase of the proportion of infectious pigs. Again, this proportion is the highest in weaned piglets, and shows a similar shape for non-weaned piglets but with smaller amplitude: some of these latter piglets remain protected by maternal antibodies. The variations of the proportion of viremic fattening pigs differ from what is observed with a constant proportion of infectious vectors: the highest proportion is almost equivalent to what is seen with weaned piglets: the majority of these fattening pigs was immunologically naïve and exposed when the proportion of infectious vectors increased. After the first peak of July the proportion of infected vectors slightly decreased but showed a second peak at the beginning of the following year: a decrease of vector population size was modeled by a decrease of the number of emerging susceptible mosquitoes, whereas the lifespan of infected vectors remained constant. The average age of the vector population thus increased when the population size decreased, which explains the increase (and the second peak) of the proportion of infected vectors. The same picture is depicted for the semi-commercial farm with slight oscillations around means explained by reproduction synchronization (Fig. 2D)

**Fig. 2.**
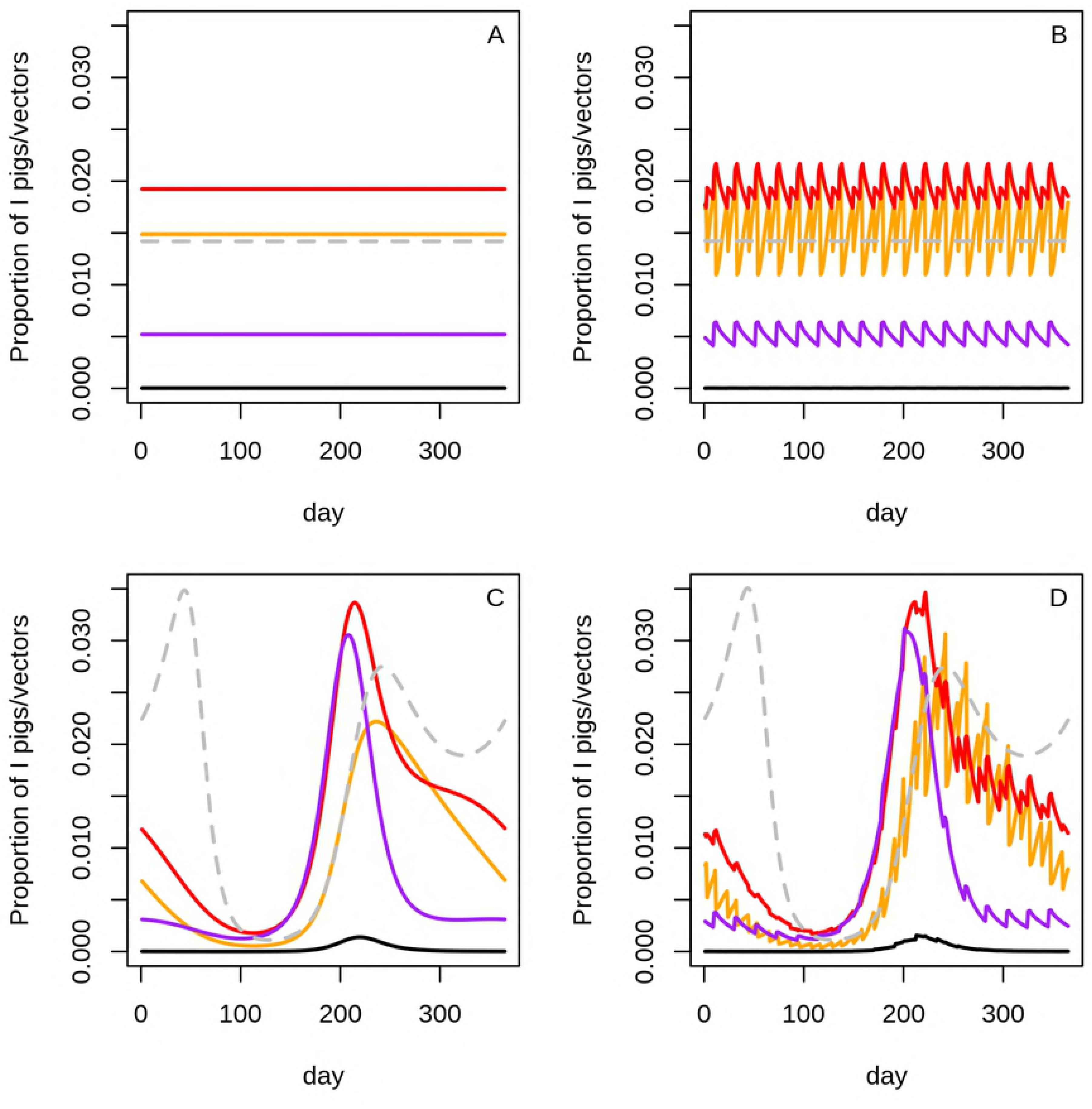
Time variations of the proportion of viremic sows (black), viremic non-weaned piglets (orange), viremic weaned piglets (red), viremic fattening pigs (purple) and infectious vectors (dashed grey), in a village breeding unit or a semi-industrial farm with constant vector population size (A and B), or in the case of seasonal vector population dynamic (C and D)

### Systematic exploration of parameter space

Fig 3, S2, S3 and S4 Fig provide the variations of the five computed output variables according to the efficacy of vector control and the sow batch interval, when the vector population size is constant and without gilt vaccination (subgraphs A), when the vector size is constant and with gilt vaccination (subgraphs B), when the vector population is seasonal and without vaccination (subgraphs C), when the vector population is seasonal and with gilt vaccination (subgraphs D).

**Fig. 3.**
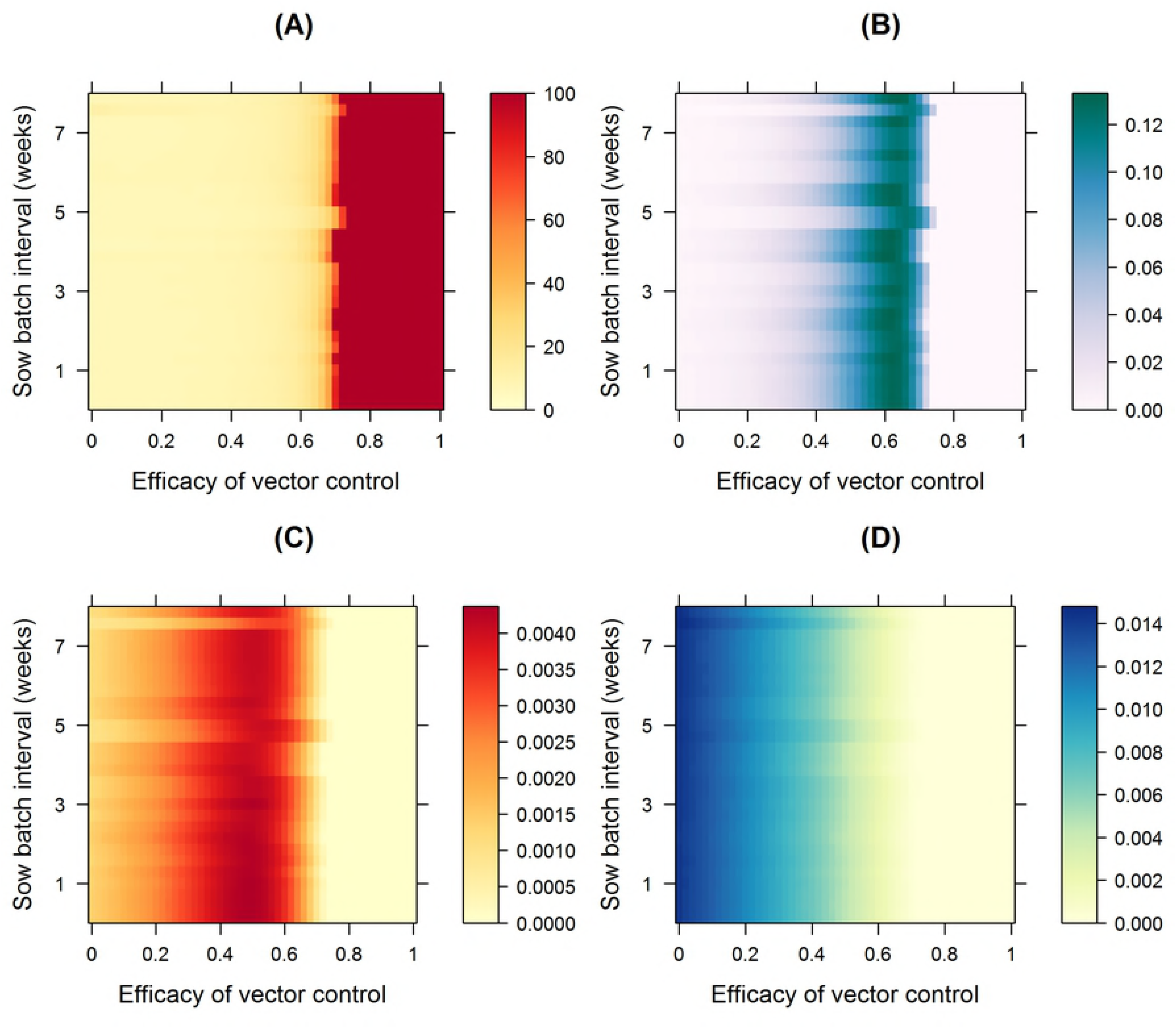
Variations at the epidemiological equilibrium, of (A) the maximal number of sows compatible with epidemic die-out, (B) the abortion rate in gilts, (C) the average proportion of viremic pigs sent to the abattoir, and (D) the average proportion of infectious mosquitoes, according to the efficacy of vector control and to the time interval between insemination of sow batches, and when no vaccination is used in gilts, in an area where the vector population size is constant.

The five computed output variables were slightly affected by variations of the time interval between insemination (Δ_*Batch*_), with and without vaccination in gilts, and the vector population size being seasonal or not. Conversely, the efficacy of vector control (*u*) has a major effect on these output variables (Fig. 3, S2 Fig, S3 Fig, S4 Fig).

The maximal number of sows per batch compatible with epidemic die-out was low when the vector population was constant (Fig. 3, S2 Fig, subgraph A) and when the efficacy of vector control was weak. Figure 4 shows the evolution of computed outputs variables in the case of a semi-commercial farm, and an interval between insemination of 21 days (i.e. corresponding to a y-value of 3 in Fig. 3): in this case, epidemic die-out could not be obtained when the number of sows per batch was >10 (Fig. 4, subgraph A). Epidemic die-out became however possible with a large number of sows per batch when the efficacy of vector control exceeded a given threshold. This threshold varied only slightly according Δ_*Batch*_ to (Fig. 3, S2 Fig, subgraph A), and was about 0.7 in a semi-industrial farm (Fig. 4, subgraph A).

**Fig. 4.**
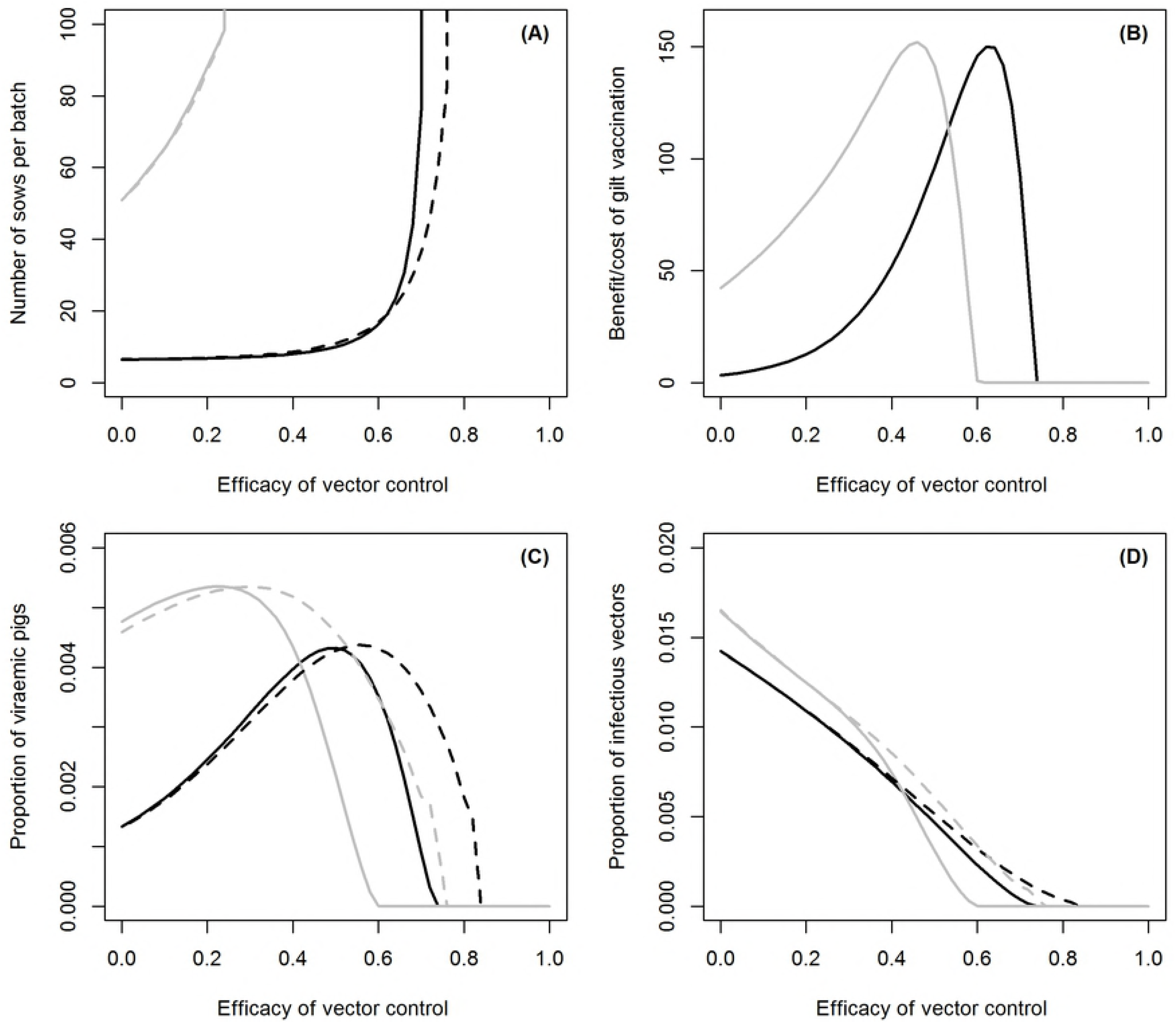
In a semi-industrial farm with a insemination interval of 21 days, and at the epidemiological equilibrium, variations of (A) the maximal number of sows compatible with epidemic die-out, (B) the benefit/cost ratio of vaccinating gilts, (C) the average proportion of viremic pigs sent to the abattoir and, (D) the proportion of infectious vectors, according to the efficacy of vector control, when gilts are vaccinated before the first insemination (dashed lines) or not (plain lines), in an area where the vector population dynamic is seasonal (grey) or not (black).

In an area where the vector population size was seasonal, epidemic die-out was possible with larger sow batches (S3 Fig and S4 Fig, subgraph A), this latter indicator increasing smoothly with the efficacy of vector control, from 50 sows per batch without vector control (*u* = 0) to >100 when *u* > 0.20 (Fig.4, subgraph A). The maximal number of sows per batch allowing epidemic die-out was poorly affected by the use of vaccination in gilts.

The abortion rate in gilts was only slightly affected by Δ_Batch_, and showed a more complex pattern when the efficacy of vector control increased (Figs 3, and S3 Fig, subgraph B), peaking when the efficacy of vector control was 0.6 in the absence of seasonality of vector abundance (and at *u* = 0.5 when vector abundance was seasonal). At this peak, the abortion rate was significant, as abortion affected >10% of gilts. The benefit-cost ratio of vaccinating gilts (S2 Fig and S4 Fig, subgraph B) showed the same pattern. In the case of a semi-industrial farm (Fig. 4B), at the peak of this benefit/cost ratio, vaccinating gilts was profitable if the unitary cost of vaccination was 150 time greater than the net margin of the breeder per kg of carcass. It is worth noting that, although vaccinating gilts was clearly not profitable in the absence of vector control when the vector population size was not seasonal, this vaccination is profitable when vector abundance was seasonal provided that the unitary cost of vaccination was less than 50 times the net margin of the breeder per kg of carcass (Fig. 4B).

As the preceding indicators, the average proportion of viremic pigs sent to the abattoir was only slightly affected by Δ_*Batch*_, but also showed a non-linear pattern when the efficacy of vector control increased (Fig 3, S2, S3, and S4, subgraph C). In a semi-industrial farm, when no vaccination was used in gilts and in the absence of seasonality in vector abundance, this proportion of viremic pigs among those sent to the abattoir was approximately 0.1% in the absence of vector control (*u* = 0), and increased when the efficacy of vector control increased, peaking for *u* = 0.5 before decreasing to zero for *u* = 0.75(Fig 4C). The peak was reached for a higher value *u* when gilts were vaccinated (*u* = 0.6). Both patterns were left-shifted when vector abundance was seasonal.

The average proportion of infectious vectors was poorly affected by the use of vaccine in gilts, Δ_*Batch*_, and the seasonality of vector population dynamic (Fig 3, S2, S3, and S4 Figs, subgraph D). This indicator decreased almost linearly when the efficacy of vector control increased, and reached 0 for *u* > 0.80 in the case of a semi-commercial farm (Fig. 4D).

### Sensitivity analysis

The influence of model parameters on output variables was very close in a village breeding unit (Fig. 5) and in a semi-industrial farm (S5 Fig), for each of the five output variables.

**Fig. 5.**
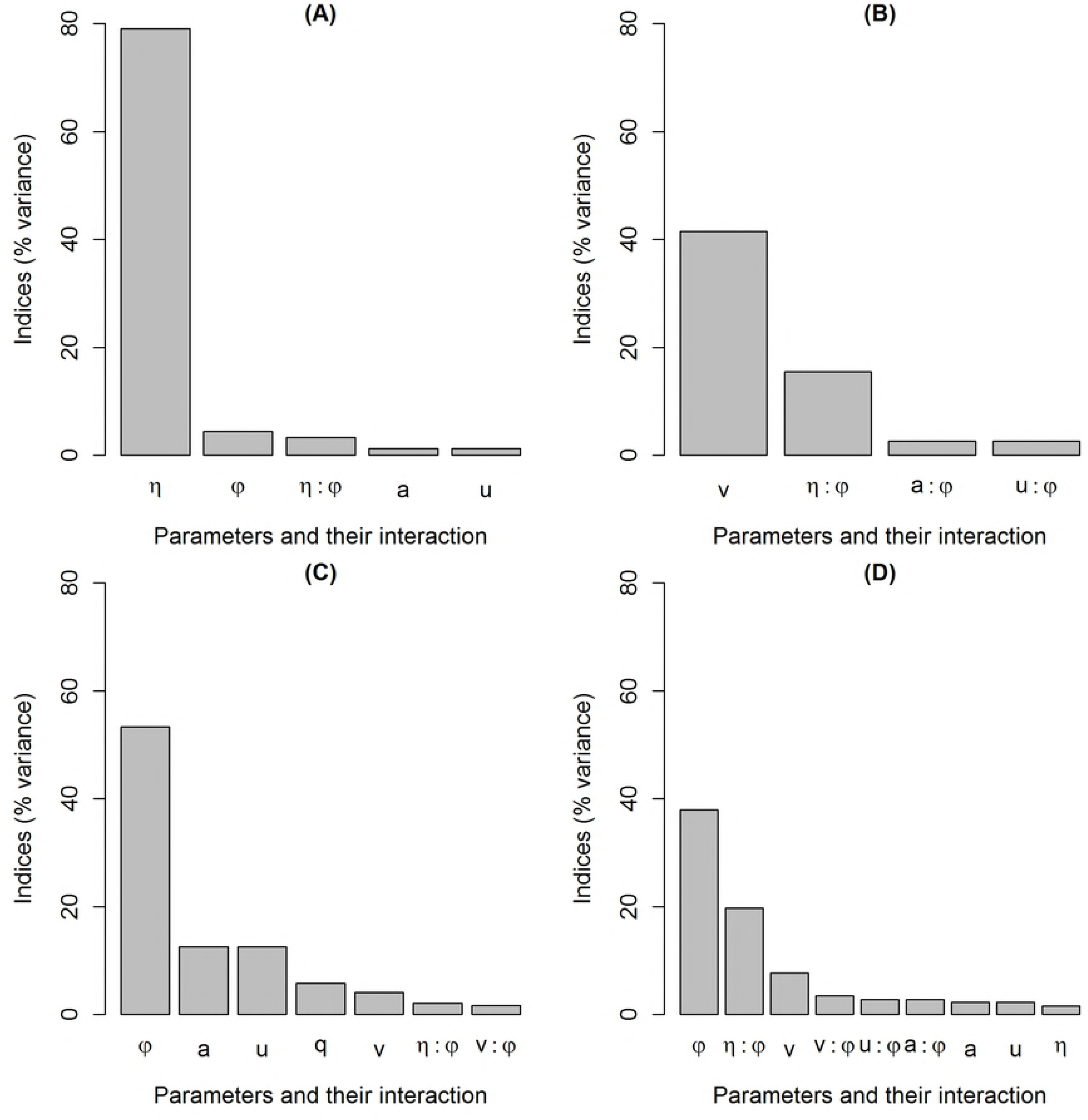
Influence of model parameters in the case of a village breeding unit, on (A) the maximal number of sows compatible with epidemic die-out, (B) the abortion rate in gilts, (C) the average proportion of infectious mosquitoes, and (D) the average proportion of viremic animals among the pigs sent to the abattoir. The y-axis represents the percentage of the total variance explained by the variation of the parameters alone, or the interaction of two parameters. The x-axis represents the parameters and their interaction. The parameters are ranked in decreasing order of the sensitivity indices, i.e. from the most to the least influential. Parameters responsible for less than 1% of the global variance were discarded from the graph.

For both farming systems, the vector population size (η) was the parameter that mostly influenced the maximal number of sows allowing epidemic die-out (Fig. 5A, S5 Fig. A). Variation of this parameter generated about 80% of the total variance, whereas all the other parameters contributed less than 10% of the total variance. The abortion rate in gilts (Fig. 5B, S5 Fig. B) was mostly influenced by vaccination (*v*), which generated about 40% of the total variance. The interaction between the vector population size (η) and the vector to host ratio (*φ*), contributed a significant part of total variance (15%). The influence of other parameters was much lower. The average proportion of infectious mosquitoes was mostly driven by the vector to host ratio (*φ*), which generated more than 50% of the total variance (Fig. 5C, S5 Fig. C). The average daily biting rate per mosquito (*a*), and the efficacy of vector control (*u*), contributed 13% of total variance. The contribution of other parameters to total variance was less than 10%. Lastly, the average proportion of viremic pigs sent to the abattoir was predominantly influenced by the vector to host ratio (*φ*) that represented about 40% of the total variance (Fig. 5D, S5 Fig. D). Again the interaction between this parameter and the vector population size (η) contributed a significant part of the total variance (20%). The contribution of other parameters was less than 10%.

## Discussion

Despite existence of vaccines, JE remains a prominent public health problem in SE Asia. With development of semi-commercial systems of pig production in rural and peri-urban areas, growing populations and increasing land use for rice-field cultures, JE threat will likely increase in the coming years in most Southeast Asian countries (2, 75). This modelling study, applied for both backyard and semi-commercial pig breeding systems, shows that combining sow vaccination and vector control could be an alternative and/or an additional measure to human vaccination to efficiently reduce both JE incidence in humans and the economic impact of JE infection on pig breeding.

Thanks to its generic structure, our model allowed representing a continuum of plausible breeding systems in Southeast Asia, ranging from a village breeding unit without synchronization but with daily births until a semi commercial farms with an interval between insemination of 8 weeks, exposed to a constant or seasonal vector pressure. However, we did not represent large commercial systems that we assumed protected by efficient biosecurity measures. This latter assumption needs to be verified. Indeed, even if the number of intensive farms remains low compared to the number of backyard and semi-commercial farms, the huge number of pigs they contain may play a role in JE transmission or persistence in a given area, and represents a threat for farm and abattoir workers, as well as people living in their close vicinity.

To assess the joint effects of the three control measures of concern, we computed five indicators: (i) the proportion of viremic vectors and (ii) the proportion of viremic pigs sent to the abattoirs, both considered as proxy of the risk of human infection; (iii) the maximum size of sow batch compatible with epidemic die out, (iv) the abortion rate of sows, and (v) the benefit-cost ratio of vaccinating gilts, related to the economic impact of JE on pig farms as well as their sustainability in an endemic context.

As expected, vector control alone is a major control tool: the maximal number of sows per batch allowing epidemic die-out increased with vector control, with constant or seasonal vector abundance, with or without gilt vaccination, and whatever the interval between births. Similarly, the average proportion of infectious vectors decreased when vector control increased. As a matter of fact, some farmers already empirically use mosquito nets – cheap, and environmentally-friendly, to protect themselves and their pigs against mosquito bites. At the village level, JEV circulation and incidence in both human and pigs could thus be largely reduced by mosquito nets use. In this work we assessed efficacy of vector control on the intensity of transmission between pigs and mosquitoes, thus indirectly on the risk of human infection. Obviously, and at least, use of insecticide-treated nets/long-lasting insecticidal nets and other personal protection methods can contribute protect humans. However recent entomological surveys performed in Cambodia showed that JE vectors are active as early as 6 pm, when people are still active (S. Boyer, pers. com; S6 Letter). Mosquito nets are therefore not sufficient to prevent human from being infected. Use of adulticides and larvicides may thus be an alternative in highly endemic areas. But both methods are costly and in the case of insecticide use, can have negative environmental consequences on non-target species that needs to be carefully assessed. Finally, integrated management approach, including alternation of wet and dry irrigation of rice fields, larval control, personal protection and improved housing and sanitation, may reduce significantly human exposure (75). The second limitation of vector control tool is its predicted paradoxical effect. When vector control is weak, infection pressure is so high that pigs get infected early in their life. When sent to the abattoir all are already immune. Gilts are also immune when covered or inseminated. There is no need to use vaccination in this case. When vector control increases until a given threshold, infection pressure decreases and piglets consequently get infected, on average, when they are older, and some may arrive at the abattoir being viremic. Similarly, a given proportion of young sows may get infected during their first gestation and abort: the benefit –cost of vaccination increases. Above this threshold that varies according to the seasonality of vector abundance, JEV transmission is interrupted or strongly reduced, and vaccination becomes again non-profitable. To be efficient, vector control level thus needs to exceed a given threshold that may be tricky to adjust under real conditions. Below this threshold, vector control may exacerbate JEV transmission, and the risk for human, even when gilts are vaccinated.

The five computed indicators of the effect of control measures on JE infection impact were only slightly affected by variations of the time interval between insemination (Δ_*Batch*_), both with and without vaccination in gilts, and the vector population size being seasonal or not. According to our results, the way this interval is managed is not an efficient tool to control JE viral circulation within pig populations. Lastly, gilt vaccination is used in temperate or sub-tropical countries, such as Taiwan or Japan. A recent survey in Vietnam showed that vaccination of future breeder pigs in epidemic areas could protect them from JE-associated reproductive disorders (31). Our modelling results corroborate this statement: gilt vaccination can be profitable to breeders of semi-industrial farms if the level of vector control is low to moderate, especially in areas where vector dynamic is seasonal.

According to the sensitivity analysis results, model parameters had the same influence on model outputs in both breeding systems. In Diallo et al, the most influential parameter driving variations of the basic reproduction number (R_0_) of two JE transmission models –one with vector borne transmission only and one incorporating both vector-borne and direct transmission, was the mosquito population size (61). Results of the sensitivity analyses in the present work are consistent with that: for both farming systems, the vector population size was the parameter that mostly influenced the maximal number of sows allowing epidemic die-out. Vector population size or/and the vector host ratio were again the most influential parameters for the proportion of infectious vectors and the proportion of viremic pigs sent to the abattoir. As expected, abortion rate in gilts is mainly driven by the vaccination rate. The direct transmission parameter, β, had a negligible influence on output variables.

The main limitation of our model is related to the entomological component. JEV is transmitted by at least 25 mosquito species (6). However, there are few data regarding JE vectors dynamics in South East Asia. In Cambodia, a recent survey performed in the Kandal province, 99% of the mosquitoes caught were JE vectors, namely *Cx tritaenhiorhynchus, Cx gelidus, Cx. vishnui* and *Cx quinquefasciatus* (47). The first three ones are considered main JE vectors in SEA (76–78). Authors of this publication showed an “apparent peak of mosquito’s abundance in May, July and December” (47). However, this trapping work was limited in time and space and obviously does not represent the dynamic of JE vectors in SEA. A second trapping survey showed a high relative abundance of these JE vectors throughout the year with a significant seasonal peak at the end of the rainy season, i.e. in October (S. Boyer, pers.com; S6 Letter). In the present work and in absence of accurate dynamic data, JE vector population was represented by a non-specific *Culex* population. Due to this lack of specificity, we chose to explore the model behavior for both constant and seasonal vector population dynamics, to take into account all plausible scenarios, and with a peak at the end of July based on Cambodian observations. This preliminary and theoretical approach needs to be refined with country, or even local-specific entomological data to take into account complex seasonality patterns and potential seasonal viral extinction that may occur in temperate sub-regions such as Northern Vietnam for instance. Secondly, and in absence of Southeast Asian data, pig breeding parameters, such as litter size or time spent in nursery, were extracted from French reports, and may not be absolutely consistent with SEA pig performances. The sensitivity analyses showed that litter size had few or no influence on the model outputs. Other parameters of the model that set the time ranges of sow production cycle and piglets grow steps, are strongly linked to pig biology. Nevertheless, gestation and suckling duration may be slightly different is SEA than in Europe. A modification of these durations would modify the interval between birth, (Δ_*Batch*_), that have no influence on model outputs. Similarly the duration between birth and slaughtering may vary from one country to another one. If this duration increases, and, in the case of a sufficient vector control, the proportion of pigs sent to the abattoir will decrease. If vector control is weak, pigs would get infected quickly: keeping pigs longer would mean keeping immune pigs. Again, the proportion of viremic pigs sent to the abattoir would decrease. On the contrary, and to answer to a higher market demand for instance, breeders may decide to sell their pigs earlier, which would increase the proportion of viremic pigs sent to the abattoir, thus increase the risk for abattoir workers and people living around to get infected.

Lastly, domestic birds may be involved in JEV transmission processes. However, and in absence of clear evidence and data, we did not incorporate any domestic bird compartment in our model. This additional compartment would have probably no influence in the case of semi- commercial farm provided that this farm breeds only pigs, but may be important for the village breeding scenario since domestic birds are present almost everywhere in SEA rural villages, and may serve as secondary amplifying and/or reservoir hosts.

As a conclusion, this preliminary work carried at the farm or village scale, demonstrates that alternative control measures can help reducing the impact of JE, both on human and pig health. Upscale this model connecting breeding systems with each other through pig movements and vector borne FOI and assess the joined effects of the three control measures of concern is the next step toward a better control of JE in Southeast Asia.

## Acknowledgments

We gratefully thank Dr Nicolas Rose (ANSES Ploufragan), Dr S. Boyer and Dr P. Dussart (Institut Pasteur du Cambodge) for their collaboration and advices.

## Supporting Information

**S1.** Letter. Personal communication P.Dussart.

**S2 Fig.** Variations at the epidemiological equilibrium, of (A) the maximal number of sows compatible with epidemic die-out, (B) benefit/cost ratio of vaccinating gilts, (C) the average proportion of viremic pigs sent to the abattoir, and (D) the average proportion of infectious mosquitoes, according to the efficacy of vector control and to the time interval between insemination of sow batches, when gilts are vaccinated before the first insemination, in an area where the vector population size is constant.

**S3 Fig.** Variations, at the epidemiological equilibrium, of (A) the maximal number of sows compatible with epidemic die-out, (B) the abortion rate in gilts, (C) the average proportion of viremic pigs sent to the abattoir, and (D) the average proportion of infectious mosquitoes, according to the efficacy of vector control and to the time interval between insemination of sow batches, when no vaccination is used in gilts, in an area where the vector population size is seasonal.

**S4 Fig.** Variations, at the epidemiological equilibrium, of (A) the maximal number of sows compatible with epidemic die-out, (B) the benefit/cost ratio of vaccinating gilts, (C) the average proportion of viremic pigs sent to the abattoir and (D), the average proportion of infectious mosquitoes, according to the efficacy of vector control and to the time interval between insemination of sow batches, when gilts are vaccinated before the first insemination, in an area where the vector population dynamic is seasonal.

**S5 Fig.** Influence of model parameters in the case of a semi-industrial farm, on (A) the maximal number of sows compatible with epidemic die-out, (B) the abortion rate in gilts, (C) the average proportion of infectious mosquitoes, and (D) the average proportion of viremic pigs sent to the abattoir. The y-axis represents the percentage of the total variance explained by the variation of the parameters alone, or the interaction of two parameters. The x-axis represents the parameters and their interaction. The parameters are ranked in decreasing order of the sensitivity indices, i.e. from the most to the least influential. Parameters responsible for less than 1% of the global variance were discarded from the graph.

**S6.** Letter. Personal communication S. Boyer.

